# Revisiting a GWAS peak in *Arabidopsis thaliana* reveals possible confounding by genetic heterogeneity

**DOI:** 10.1101/2021.02.03.429533

**Authors:** Eriko Sasaki, Thomas Köcher, Danièle L Filiault, Magnus Nordborg

## Abstract

Genome-wide association studies (GWAS) have become a standard approach for exploring the genetic basis of phenotypic variation. However, correlation is not causation, and only a tiny fraction of all associations have been experimentally confirmed. One practical problem is that a peak of association does not always pinpoint a causal gene, but may instead be tagging multiple causal variants. In this study, we reanalyze a previously reported peak associated with flowering time traits in Swedish in *Arabidopsis thaliana*. The peak appeared to pinpoint the *AOP2/AOP3* cluster of glucosinolate biosynthesis genes, which is known to be responsible for natural variation in herbivore resistance. Here we propose an alternative hypothesis, by demonstrating that the *AOP2/AOP3* flowering association can be wholly accounted for by allelic variation in two flanking genes with clear roles in regulating flowering: *NDX1*, a regulator of the main flowering time controller *FLC*, and *GA1*, which plays a central role in gibberellin synthesis and is required for flowering under some conditions. In other words, we propose that the *AOP2/AOP3* flowering-time association is yet another example of a spurious, “synthetic” association, arising from trying to fit a single-locus model in the presence of two statistically associated causative loci.

## Introduction

Genome-wide association studies (GWAS) have become an essential tool for studying the genetics of natural variation. In addition to its tremendous impact on human genetics, GWAS is being applied routinely to a wide range of species, and massive numbers of genotype-phenotype associations have been revealed (Atwell *et al*., 2010; Flint and Eskin, 2012; MacArthur *et al*., 2017). However, only for a tiny subset do we have any idea of why the association exists, *i*.*e*., the molecular mechanism. There are many reasons for this, but an important one is that peaks do not always pinpoint the causal genes (Hormozdiari *et al*., 2014; Tam *et al*., 2019). In settings where the environment cannot be controlled, spurious associations may simply arise because of environmental confounding, but our focus here is on confounding by the genetic background, and in particular on genetic background effects that are not sufficiently diffuse to be readily be removed by approximate methods like kinship or Principal Component corrections (Atwell *et al*., 2010; Vilhjálmsson and Nordborg, 2013). Such effects may arise whenever there is linkage disequilibrium between non-trivial allelic effects (Platt *et al*., 2010; Dickson *et al*., 2010), and is a well-known problem for fine-mapping when there is *allelic* heterogeneity. In *Arabidopsis*, examples include multiple functional alleles of *FRIGIDA* (*FRI*) *(Atwell et al*., *2010)*, and *DELAY OF GERMINATION1 (DOG1)* (Kerdaffrec *et al*., 2016). Less clear is how frequently spurious associations arise from *genetic* heterogeneity, *i*.*e*., from alleles for different genes affecting the same trait. In this paper we discuss what we believe to be an example of this: causal allelic variation at two different genes, separated by roughly 120 kb, inducing a spurious peak of association in a third gene located between the causal loci.

## Results

### GWAS for flowering time suggested a role for *AOP2*

Flowering time is an adaptively and agriculturally important trait that has been intensively studied in *Arabidopsis thaliana*. Thanks to decades of functional work, the pathways involved in flowering time regulation (and their interaction with the environment) are extremely well understood (Koornneef *et al*., 1998; Srikanth and Schmid, 2011; Andrés and Coupland, 2012). Known flowering-time regulators are also highly variable in nature, and GWAS for flowering time typical reveal a variety of known loci (Brachi *et al*., 2010; Atwell *et al*., 2010; Li *et al*., 2010; Sasaki *et al*., 2015; 1001 Genomes Consortium, 2016; Zan and Carlborg, 2019)

A major source of natural variation for flowering is a variety of loss-of-function alleles of *FRI*, which regulates *FLOWERING LOCUS C* (*FLC*), a key regulator of flowering time, crucial for helping plants flower in the right season by “remembering” exposure to cold winter temperatures using a fascinating epigenetic mechanism (Whittaker and Dean, 2017). Indeed, *FRI* was first identified through natural variation (Johanson *et al*., 2000), but, despite explaining a considerable fraction of the variation for flowering time, has proven difficult to map using GWAS, mostly because it has such high allelic heterogeneity (Atwell *et al*., 2010). However, several GWAS identified a strong peak roughly 1 Mb from *FRI*, a peak which stood out because it was not obviously associated with a known flowering time gene (Brachi *et al*., 2010; Atwell *et al*., 2010; Li *et al*., 2010; Zan and Carlborg, 2019). Instead, this peak appeared to pinpoint the highly variable and adaptively important *ALKENYL HYDROXALKYL PRODUCING* (*AOP*) cluster (Atwell *et al*., 2010; Kerwin *et al*., 2011; Katz *et al*., 2020), containing three tandemly duplicated genes involved in the synthesis of glucosinolates, secondary metabolites that play important role in defense against herbivory (Kliebenstein *et al*., 2001). In the Swedish population, the strongest association was found for the SNP at chr4:1355057, 961 bp upstream of *AOP2*, and strongly correlated both with flowering time and *FLC* expression (Figs. 1 and S1). Here we focus on this association, and return to associations in other populations in the discussion.

**Fig. 1.**
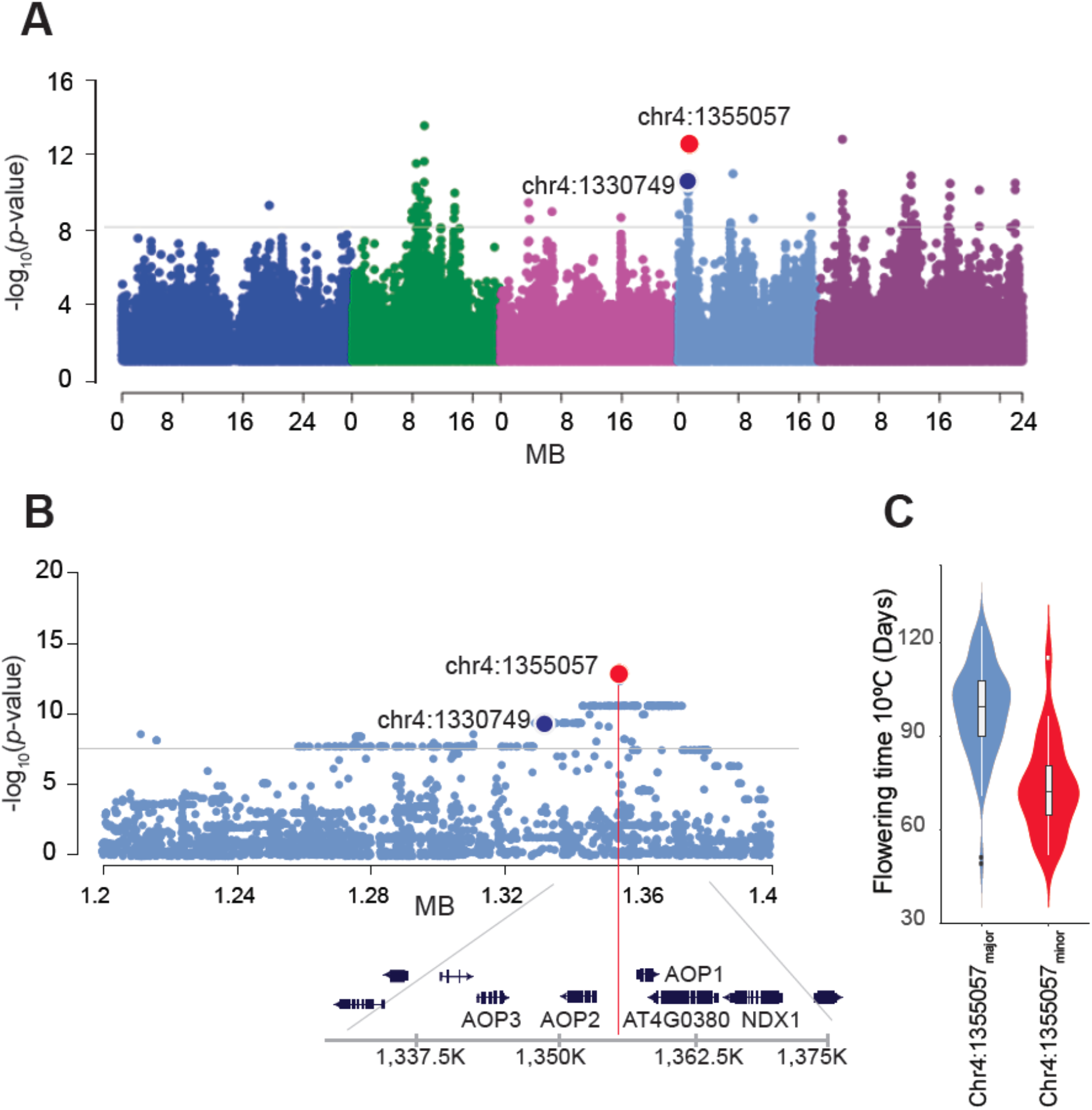
GWAS for flowering time revealed a peak centered on the chromosome 4 *AOP* cluster. (A) Genome-wide Manhattan plot for flowering time at 10°C in 132 Swedish lines, with SNP reported in Atwell et al. (2010) highlighted in blue (chr4: 1330749), and the strongest association in red (chr4: 1355057). A linear model without structure correction was used (see Methods). (B) Zoom-in on the peak, with gene annotation. (C) Violin plot showing the difference in flowering time between lines carrying major and minor alleles at chr4:1355057.

### Functional variation at *AOP2* does not affect flowering time

Although a transgene experiment had shown that *AOP2* (but not *AOP3*) could affect flowering time (Kerwin *et al*., 2011; Jensen *et al*., 2015), pleiotropic effects on flowering time are common (Chong and Stinchcombe, 2019), and we were not convinced that this was the explanation for the *AOP2* peak. To investigate this further, we first explored the functional *AOP2* variants tagged by chr4:1355057. Jensen et al. (2015) showed that *A. lyrata AOP2* delays flowering when overexpressed in the reference line Col-0, which carries a natural non-functional *AOP2* allele due to a 5-bp deletion causing a frameshift and leading to accumulation of different glucosinolates (Kliebenstein *et al*., 2001). They suggested that delayed flowering results from an interaction between the glucosinolate and flowering pathways. Based on these results, functional *AOP2* alleles should be associated with delayed flowering in *A. thaliana. AOP2* has multiple alleles inducing frameshift in addition to the Col-0 allele (Neal *et al*., 2010). In the Swedish population, five indels that could induce frameshift were identified, including the Col-0 type (Fig. 2A; Table S1).

**Fig. 2.**
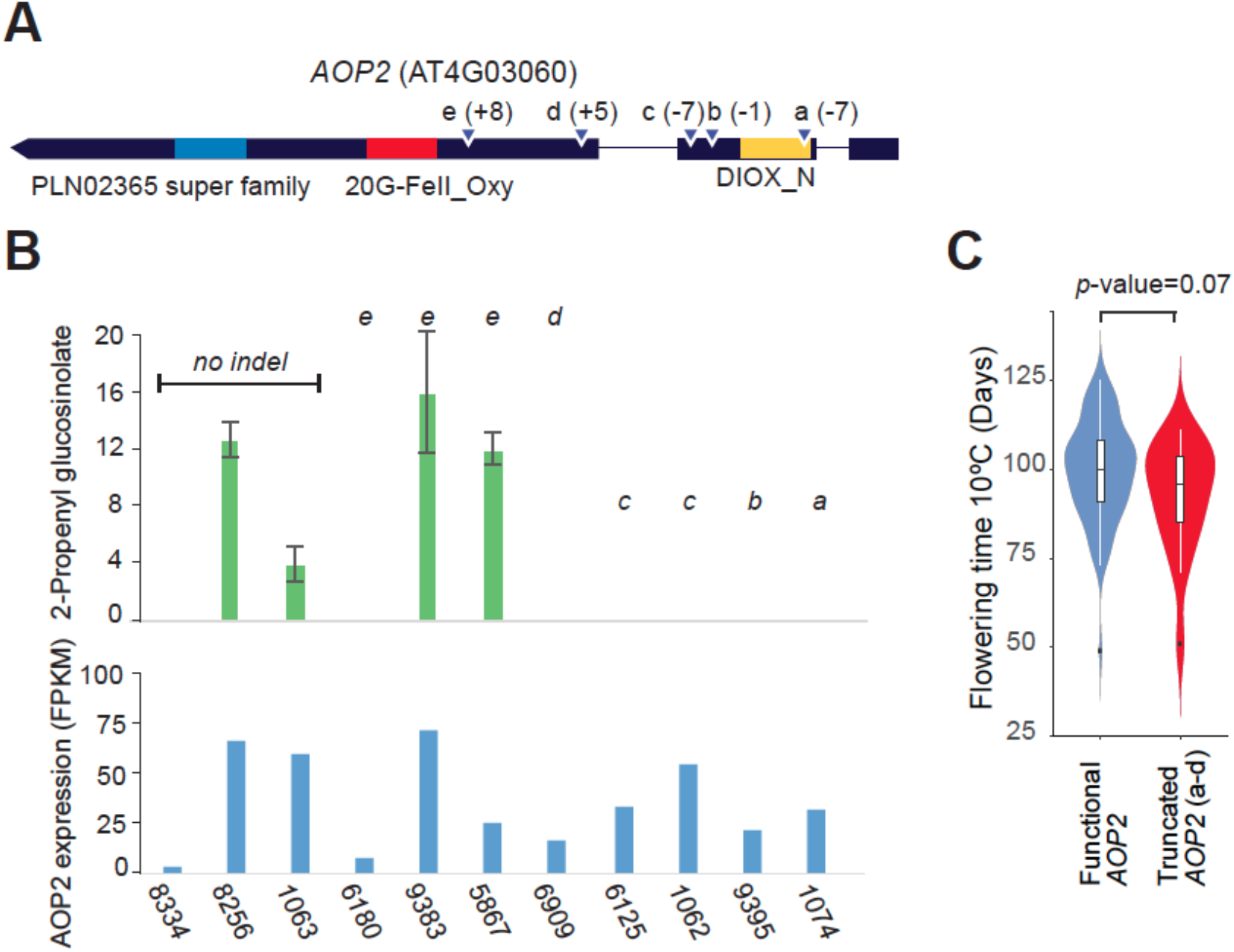
Functional validation of *AOP2* alleles. (A) The gene model of AOP2.1. Colored parts are conserved protein domains. Triangles show predicted indels that could cause frameshift. + and - are the indel size from the reference sequence. (B) Accumulation of 2-propenyl glucosinolate in each type of allele and the *AOP2* expression. Marks from a to e are corresponding to the indels. (C) The flowering time based on *AOP2* indel genotypes. Functional *AOP2* is the genotype without frameshift and truncated *AOP2* has frameshift (a-d). Only chr4:1355057 reference lines with expressed *AOP2* were used for the plots to control for the effect of chr4:1355057 (which is of course known to be associated with flowering time variation).

We assessed the functional effect of these five indels directly using mass spectrometry (Fig. 2B). AOP2 converts 3-methylsulfinylpropyl and 4-methylsulfinylbutyl glucosinolate to 2-propenyl and 3-butenyl glucosinolate, respectively (Kliebenstein *et al*., 2001). Lines having any indel in the second exon (a to c in Fig. 2A) or the Col-0 indel (d) in the third exon did not accumulate 2-propenyl and 3-butenyl glucosinolate, although transcripts were detected in all cases (Fig. 2B; Table S2). The fifth insertion (e) did not appear to affect 2-propenyl and 3-butenyl glucosinolate accumulation significantly.

Contrary to a causal role for glucosinolate pathways in flowering time variation, *AOP2* functionality is neither correlated with chr4:1355057 nor flowering time. For example, all 105 lines carrying the reference allele at chr4:1355057 show substantial *AOP2* expression, and 19 lines of them have indels that disrupt function (Table S1), but this has no significant effect on flowering time (Fig. 2C). There is a weak correlation between *AOP2* expression and flowering (Fig. S1B), but it seems unlikely that this relationship reflects causality, when functional allelic variation does not. In conclusion, although it has been demonstrated that *AOP2* can affect flowering time (Kerwin *et al*., 2011; Jensen *et al*., 2015), the considerable functional *AOP2* variation observed in the Swedish population is not strongly correlated with flowering time, suggesting that the major association between chr4:1355057 and flowering time arises for other reasons.

### The *AOP2* peak tags a diverged allele of *NDX1*

In order to identify potential causal variants, we dissected the local haplotype structure surrounding chr4:1355057 using principal component analysis (PCA) (Fig. 3A). Consistent with the fact that the Swedish population has a strong north-south population structure (Long *et al*., 2013), the first two principal components with the latitude of origin (PC1 *r*^*2*^=0.28; PC2 *r*^*2*^=0.25). However, the third principal component was not correlated with global structure, but rather identified an extended haplotype carried by 20 of the 51 lines that also carried the non-reference chr4:1355057 allele (Fig. 3A). This haplotype (denoted chr4:1355057b) contained three genes upstream of *AOP2*, including *NDX1* (AT4G03090; chr4:1366053..1371237) — a known regulator of *FLC* that binds to the promoter region of *COOLAIR*, the antisense transcript of *FLC*, and inhibits the degradation of *FLC* by stabilizing the R-loop (Sun *et al*., 2013) (Fig. 3B).

**Fig. 3.**
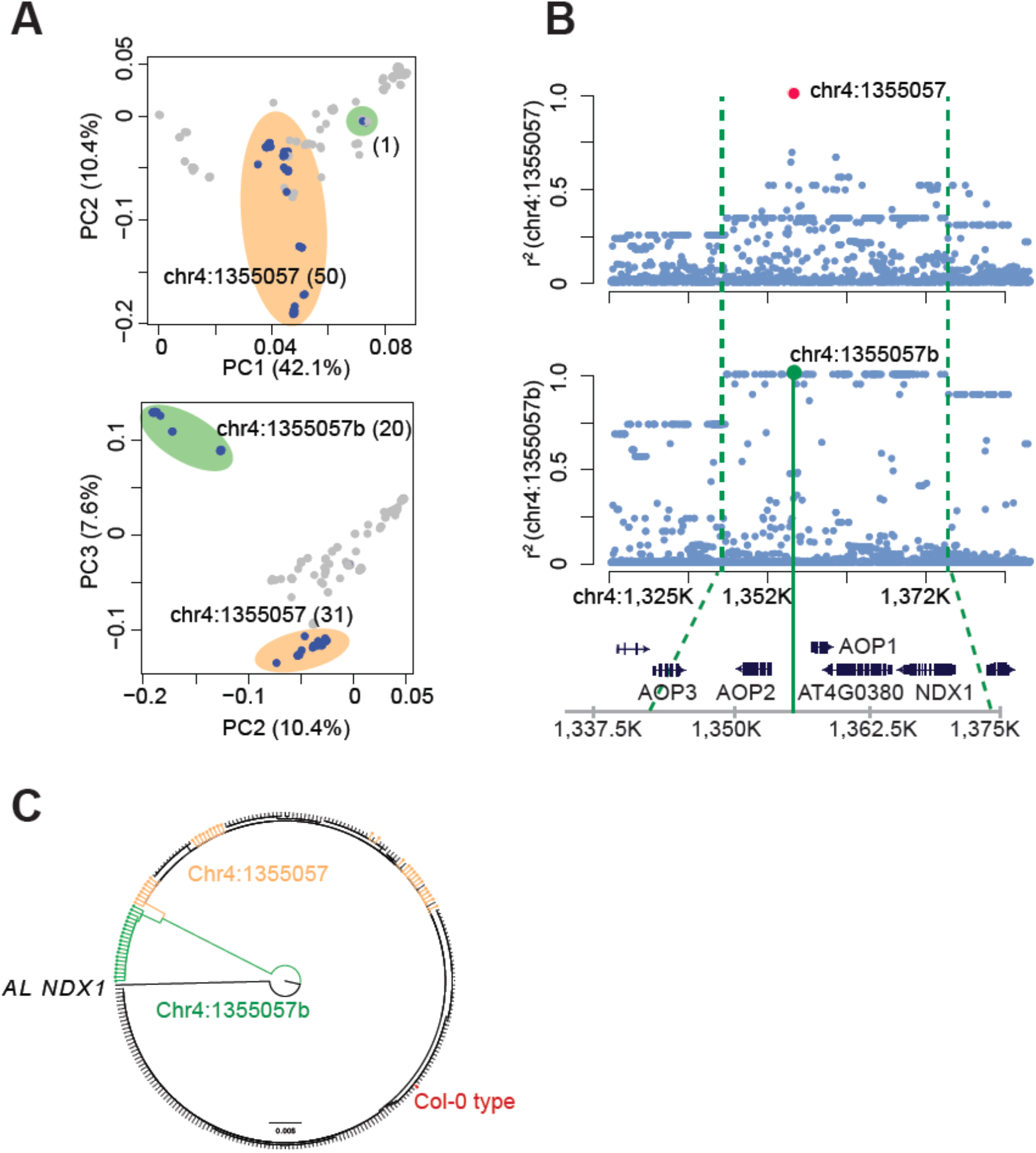
Haplotype structure around the *AOP* peak. (A) PCA of SNPs in the 60 kb region around chr4:1355057. Major (reference type) and minor alleles of chr4:1355057 are plotted in grey and blue, respectively. The clusters of minor chr4:1355057 alleles are indicated in orange and green, respectively. Numbers in brackets are allele counts. (B) Decay of linkage disequilibrium with respect to chr4:1355057 and chr4:1355057b, the subset of the chr4:1355057 minor alleles associated with an extended haplotypes (the borders of which are indicated by vertical green lines) (C) Neighbor-joining tree of *NDX1* alleles in the Swedish population (n=259). Diverged *NDX1* haplotypes associated with the chr4:1355057b haplotype are shown in green, while orange denotes remaining lines associated with rest of the chr4:1355057 minor allele.

Furthermore, the chr4:1355057b haplotype is perfectly associated with a highly diverged *NDX1* allele (Fig. 3C). The non-synonymous sequence divergence between this allele and the reference allele is close to 1%, and the changes are predicted to affect gene activity (Sun *et al*., 2013) (Fig. S2). Mutant lines confirmed that *NDX1*, unlike neighboring genes, has a significant effect on flowering (Fig. S3).

### Multilocus GWAS including *NDX1* reveals a new association near *GA1*

These observations suggested that the flowering time association peak centered on chr4:1355057 could partly be due allelic variation at *NDX1* (Fig. 4A). To explore this further, we performed GWAS while including the chr4:1355057b haplotype as a cofactor to regress out the effect of the *NDX1* polymorphism (Fig. 4B). Doing so did not eliminate the significant peak on chromosome 4 (suggesting that *NDX1* is not the only causal variant), but moved it over 100 Kbp in the opposite direction of *NDX1*. The “new” peak was quite broad and flat, but the second strongest association (chr4:1236543; −log_10_*p*-value=11.85; MAC=15) was 1.1 Kbp downstream of another well-known flowering regulator, *GIBBERELLIC ACID REQUIRING 1* (*GA1* Chr4:1237671..1244822) that is essential for gibberellic acid biosynthesis (Sun and Kamiya, 1994). Gibberellin plays a crucial role in the transition to flowering through regulation of *LEAFY* (*LFY*) and *FLOWERING TIME LOCUS T* (*FT*) (Blazquez *et al*., 1998; Porri *et al*., 2012), and loss of function mutants of *GA1* cannot flower under certain conditions (Reeves and Coupland, 2001).

**Fig. 4.**
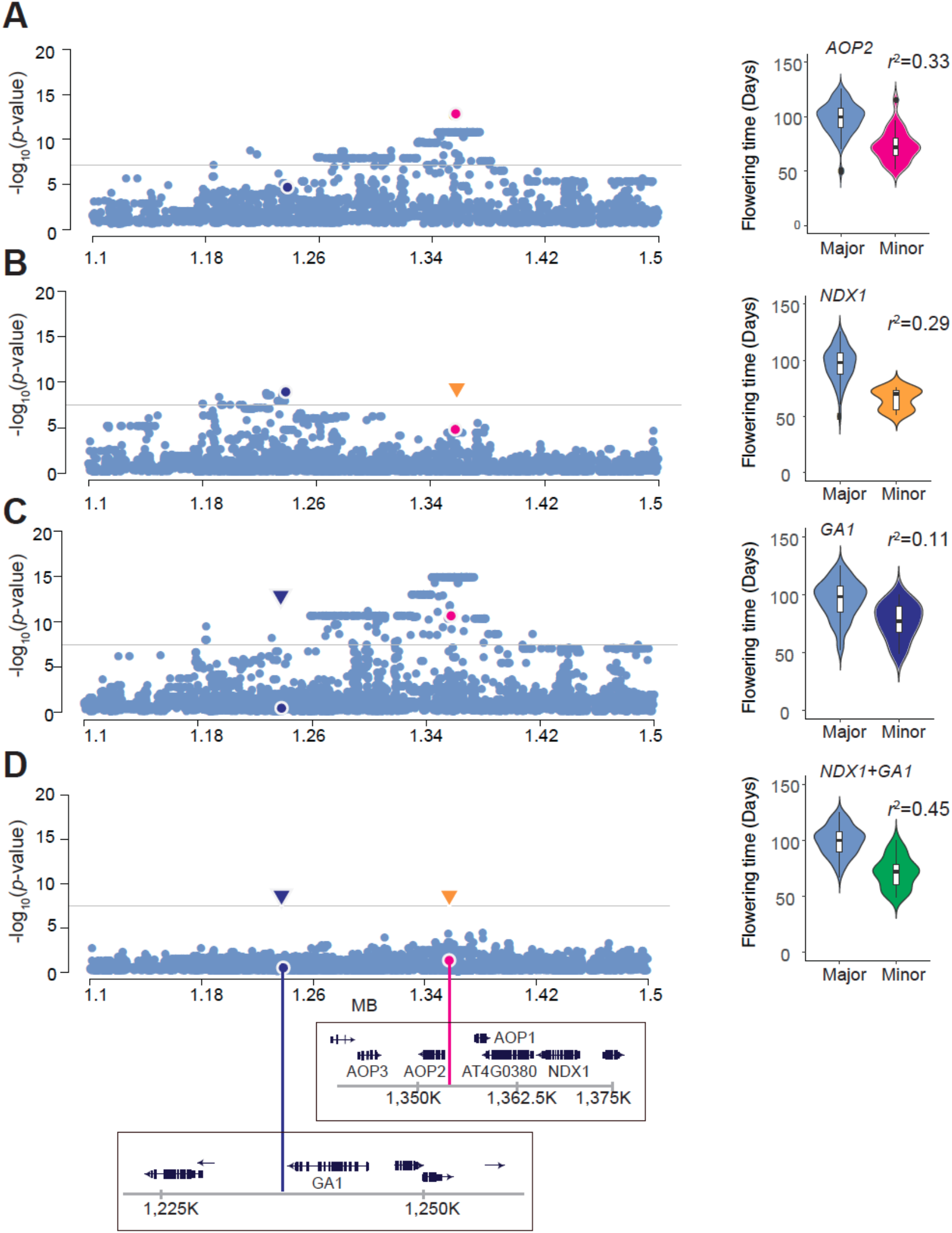
Conditional GWAS suggests genetic heterogeneity. Zoom-in plots of conditional GWAS surrounding chr4:1355057 with violin plots illustrating how much of the variation is explained by each model. Arrows on the Manhattan plots indicate SNPs used for the cofactors. (A) The original association identified an association in *AOP2* (chr4:1355057, magenta) that explained 33% of the flowering time variation. (B) Conditional GWAS using a diverged *NDX1* haplotype (chr4:1355057b, orange) that explained 29% of the variation revealed an association near *GA1* (chr4:1236543, dark blue). (C) Conditional GWAS using *GA1* peak that explained 11% of the variation. (D) Conditional GWAS using both *NDX1* and *GA1* as co-factors fully explained the original peak (explaining 45% of the variation).

Previous GWAS and linkage mapping studies have suggested that allelic variation at *GA1* plays a role in flowering time variation, and the association is known to be sensitive to population structure correction (Brachi *et al*., 2010). These kinds of problems are often caused by extensive linkage disequilibrium, the existence of which is evident (Fig. S4). This can also be seen by carrying out a third GWAS, now with the chr4:1236543 (*GA1*) SNP as a cofactor, because this causes an increase in the height of the *AOP* peak demonstrating that these peaks are indeed not independent (Fig. 4C).

### Polymorphisms at *GA1* and *NDX1* jointly explain the *AOP* association

Finally, we asked whether allelic variants at *GA1* and *NDX1* were jointly sufficient to explain the peak centered on *AOP2*. When we performed GWAS using both chr4:1236543 (*GA1*) and chr4:1355057b (*NDX1*) as cofactors, the peak at the *AOP* cluster completely disappeared (Fig. 4D) — just as if we had taken chr4:1355057 (*AOP2*) as a cofactor (Fig. S5). The distribution of phenotypes explained by chr4:1355057 is consistent with a cumulative contribution by rarer alleles at chr4:1355057b and chr4:1236543 (Fig. 4D), and explain more of the variation (as expected given the extra parameter). The same pattern was seen in GWAS for *FLC* expression (Figs. S5 and 6). These results suggest that the major flowering time association at the *AOP* cluster may be a spurious, “synthetic” association that results from the complex pattern of linkage disequilibrium between causal polymorphism at two nearby loci, *GA1* and *NDX1*. The existence of extensive linkage disequilibrium (Fig. S4) and haplotype structure (Fig. S7) in the region is clear, although we note the average decay of linkage disequilibrium is by no means unusual relative to the rest of the genome (Fig. S8).

## Discussion

In this paper we have presented an alternative interpretation for a published GWAS peak, a potential example of genetic confounding. We demonstrate that a reproducible association between flowering time (and *FLC* expression) and SNPs in the *AOP2/AOP3* glucosinolate biosynthesis cluster can alternatively be explained using a two-locus model, where the causal variants are in two flanking genes directly involved in the regulation of flowering: *GA1*, essential for gibberellin synthesis, and *NDX1*, a regulator of *FLC*. Under this interpretation, the *AOP2* peak is a spurious association (Figure 5), an artefact of incorrectly fitting a single-locus model in the presence of two causative loci (Platt *et al*., 2010; Dickson *et al*., 2010; Atwell *et al*., 2010). The problem is analogous to the problem of “ghost QTLs” in classical linkage mapping (Haley and Knott, 1992; Martínez and Curnow, 1992).

**Fig. 5.**
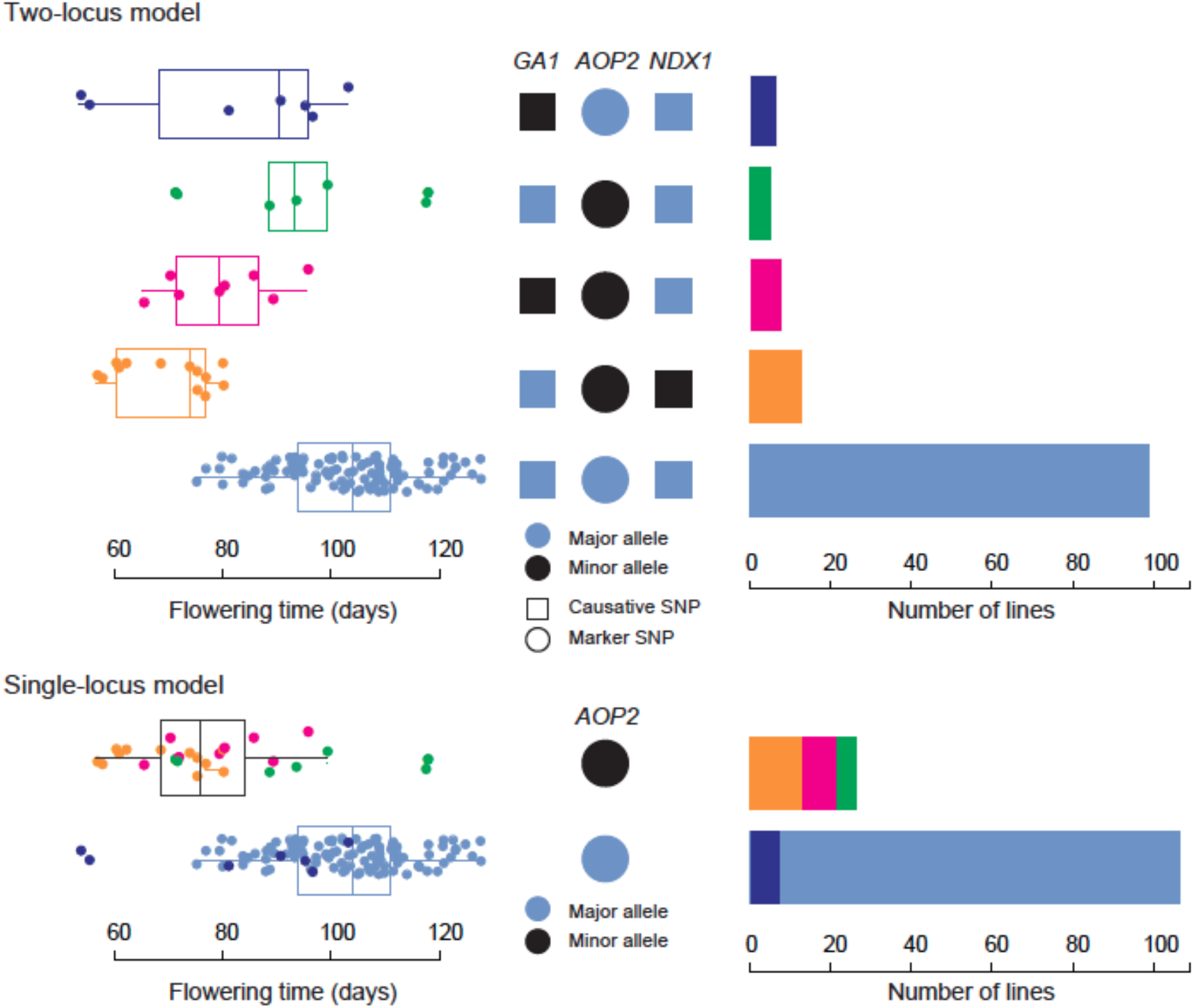
Summary of our results. We consider three di-allelic SNPs, in (or near) *GA1, AOP2*, and *NDX1*, respectively. Because of strong linkage disequilibrium, the minor *NDX1* allele is only found on haplotypes with the major *GA1* and minor *AOP2* allele, which means there are only five haplotypes — three are missing. The figure shows the flowering time distribution and observed frequency for each of these haplotypes. Our model is that the minor *GA1* and *NDX1* alleles are tagging early-flowering alleles at these two loci, and that the minor *AOP2* allele is associated with early flowering because it is the best single locus that tags both of these loci. Colors in the single-locus model correspond to those in the two-locus model.

We emphasize that neither of these two models has been experimentally confirmed. Merely knocking genes out is not sufficient for a trait like flowering time, which has been shown to be highly “omnigenic” (Boyle *et al*., 2017) in the sense that random knock-out mutations are as likely as *a priori* candidates to affect it (Chong and Stinchcombe, 2019). Because direct gene replacement is not feasible in *A. thaliana*, experimental testing of these two models would thus entail knocking out the native allele at multiple loci in multiple genetic backgrounds, and replacing it with cloned native alleles, using on the order of 50 independent transgenic lines per construct to account for position effects (Li *et al*., 2014). In late-flowering *A. thaliana*, with an essentially annual life-cycle, this would be a multi-year project requiring considerable resources.

That said, we believe that our two-locus model (involving two known flowering regulators) is a more likely explanation for the association seen here than the single-locus model (involving glucosinolate production). We say this primarily because, although flowering time is clearly “omnigenic” in the sense of presenting a large mutation target, GWAS results for flowering time (like many other phenotypes in *A. thaliana*) have generally showed a strong over-representation for genes in known pathways (Atwell *et al*., 2010; Sasaki *et al*., 2015). This is in sharp contrast to human genetics, where GWAS results have generally been extremely difficult to interpret (Boyle *et al*., 2017). A plausible explanation for this difference is that much of the variation in *A. thaliana* is adaptive (Atwell *et al*., 2010).

Indeed it may well be the case that selection is indirectly responsible for the *AOP2* flowering time association. The role of the *AOP* cluster in defense against herbivory is well established (Kliebenstein *et al*., 2001), and it is tempting to speculate that strong selection on glucosinolate variation could have contributed to the complex haplotype structure in the region — leading to associations between SNPs in *AOP2* and functional variants in nearby flowering regulators simply through random hitchhiking (Maynard Smith and Haigh, 1974). It should be noted that while a two-locus model appears to be required to explain the *AOP2* flowering-time association seen in the Swedish population, the association seen in two other samples (Atwell *et al*., 2010; Li *et al*., 2010), can simply be explained by extremely strong linkage disequilibrium between *AOP2* variants and the diverged *NDX1* haplotype described above (Figure S9). Spurious flowering time associations due to regional selection on other traits has also been suggested for maize (Larsson *et al*., 2013).

To conclude, while we may never know which (if any) of the models proposed here is correct, there is no doubt that spurious associations like this do exist, and may complicate interpretation of mapping results (Huang *et al*., 2010; Atwell *et al*., 2010; Larsson *et al*., 2013; Hormozdiari *et al*., 2014; Kerdaffrec *et al*., 2016). Although representing a difficult model-selection problem, better methods for systematically identifying such associations could be a very cost-efficient way of getting more information out of GWAS results.

## Materials and methods

### Data sets

We used published *A. thaliana* data sets containing genotypes (Long *et al*., 2013), RNA-seq transcriptome data (Dubin *et al*., 2015), as well as flowering time phenotypes (Sasaki *et al*., 2015, 2017) for the Swedish population. All plants were grown under a constant 10°C (132 lines) in 16 h day length condition. For RNA seq analysis, RNA was extracted from whole rosettes collected at 11-12 h after dawn at nine-leaf stage (Dubin *et al*., 2015). Other phenotype data, including *FLC* expression (Atwell *et al*., 2010) and flowering time under Sweden spring condition in 2008 (Li *et al*., 2010), were obtained from AraPheno (Seren *et al*., 2017).

### Plant materials

Loss-of function mutants of AT4G03080 (SALK_051383), AT4G03100 (SALK_082878), and *NDX1* (WiscDsLox344A04) (Sun *et al*., 2013) were grown with the wild type under a constant 21°C in 16 h day length condition.

### Statistical analysis

#### GWAS

GWAS was performed using LIMIX version 3.0.4 (Lippert *et al*., 2014) with full genome SNPs in a Swedish population (*n*=132) (Long *et al*., 2013) and 250K SNP chip genotypes in RegMap panel (Horton *et al*., 2012). A linear mixed model (LMM) was used to correct population structure with a kinship matrix representing genetic relatedness (IBS) (Yu *et al*., 2006; Kang *et al*., 2008). For GWAS without correction of population structure (Figs 1, 4, S1, S5, and S6), a linear regression model was used. Uncorrected GWAS was used because several of the variants in the chromosome 4 region are strongly correlated with population structure in the Swedish population, rendering fine-mapping impossible because of lack of power. Note that our primary interest is not the genome-wide significance of the region, but rather identifying potential causal SNPs within it. It has previously been observed that kinship correction can obscure causality locally (Kerdaffrec *et al*., 2016). Cofactor analysis was performed using a multiple linear regression model without population structure correction in the Swedish population (Figs 4, S1, S4, and S5), and LMM for global population (Fig S9).

#### PCA

The entire Swedish population (259 lines (Long *et al*., 2013)) were used to analyze local genetic structure around the AOP cluster. SNPs in the 60 kb region around chr4:1355057 were extracted and analyzed using the prcomp function in R (https://www.r-project.org).

#### Haplotype analysis

For the analysis, SNPs covering *GA1* and *NDX1* regions, including the 30 Kbp upstream of *GA1* and downstream of *NDX1*, were used from a pre-imputation version of the Regional Mapping Project SNP panel, including 1307 global lines (Horton *et al*., 2012). These SNPs were used as the input into fastPHASE version 1.4.8 (Scheet and Stephens, 2006), which was run using the default settings as described in (Li *et al*., 2014).

### Genotyping *NDX1* and *AOP2*

For population samples, amino acid sequences of *NDX1* and *AOP2* were predicted using genome data, including SNPs and short indels (Long *et al*., 2013). *NDX1* sequences of Col-0 and chr4:1355057b alleles were also confirmed by Sanger sequencing after cloning the 7.8 Kbp region with forward primer 5’-CTGGTAAATACTGTGTGTAGACAATTCT-3’ and reverse primer 5’-TCGATGTTTGACGGCAAAGGATGAAG-3’. Line 6180 (T□ÄL 07; latitude 62.6322 longitude 17.6906) was chosen to represent chr4:1355057b alleles. All predicted chr4:1355057b allele-specific SNPs were confirmed by the Sanger sequencing.

### Measurement of expression levels

For population samples, *FLC* and *AOP2* expressions were extracted from RNA-seq data of leaf tissue under 10°C constant temperature (Dubin *et al*., 2015). For mutants, total RNA was extracted from aerial parts of nine-leaf stage seedlings collected at 8 h after dawn using RNeasy mini kit (Qiagen) with DNase treatment (Thermo Fisher Scientific). cDNA was synthesized using the SuperScript III First-Strand Synthesis System (Invitrogen). qRT-PCR was performed using the LightCycler 96 system (Roche) with FastStart Essential DNA Green Master (Roche). *SAND* (AT2G28390) was used for a control to normalize the transcript abundance (Czechowski *et al*., 2005) using the ddCT method. The primer’s sequences were *SAND*: 5’-AACTCTATGCAGCATTTGATCCACT-3’ and 5’-TGATTGCATATCTTTATCGCCATC-3’, and *FLC*: 5’-TGAGAACAAAAGTAGCCGACAAG-3’ and 5’-ATGCGTCACAGAGAACAGAAAGC-3’.

### Assessment of AOP2’s functionality

#### Tissue disruption

Frozen 10 mg samples in liquid nitrogen in 2 ml Eppendorf tubes were stored at −80 °C. Precooled 1 ml 90% methanol (90% MeOH/10% 10 mM ammonium bicarbonate in H_2_O; −20 °C) was added by the final methanol concentration remaining above 78%. Tissue was disrupted by adding two small stainless beads bearings and agitating with a tissue lyser (tissuelyser II, Qiagen, Hilden, Germany) for 10 min at 20 rev/s. The sample was shaken for a further 60 min (70 rev/s) in the cold. After centrifuge at 13,000 rpm and the supernatant was transferred to a fresh tube and discard the pellet.

#### Measurement

Measurement of glucosinolates were performed according to a previous study (Liang *et al*., 2018). Briefly, 2 µl of each sample was injected into a SeQuant ZIC-pHILIC HPLC column (Merck, 100 × 2.1 mm; 5 µm), and the respective guard column operated with an Ultimate 3000 HPLC system (Dionex, Thermo Fisher Scientific) at a flow rate of 100 µl/min. The HPLC was directly coupled via electrospray ionization in the negative ion mode (2.8 kV) to a TSQ Quantiva mass spectrometer (Thermo Fisher Scientific). A linear gradient (A: 95% acetonitrile 5%, 10 mM aqueous ammonium acetate; B: 5 mM aqueous ammonium acetate) starting with 5% B and ramping up to 45% B in 9 minutes was used for separation. Chromatograms were interpreted using TraceFinder (Thermo Fisher Scientific) and manually validated. The following transitions were used for relative quantitation: 3-hydroxypropyl glucosinolate m/z 376 → m/z 97; m/z 376 → m/z 259, 2-propenyl glucosinolate m/z 358 → m/z 97; m/z 358 → m/z 75, 4-hydroxybutyl glucosinolate m/z 390 → m/z 259; m/z 390 → m/z 97; m/z 390 → m/z 75 and 3-(methylsulfinyl)propyl glucosinolate m/z 422 → m/z 259; m/z 422 → m/z 97; m/z 422 → m/z 75.

## Acknowledgments

The authors would like to thank Caroline Dean for sharing knowledge and seeds of the *AtNDX1* mutant. We also thank Ümit Selen for technical support with the data analysis, and Daniel J. Kliebenstein and Haijun Liu for comments and helpful discussions. The VBCF Metabolomics Facility is supported by the City of Vienna through the Vienna Business Agency.

## Conflict of Interest

The authors have no conflict of interest to declare.

## Notes

### Competing Interest Statement

The authors have declared no competing interest.

