## Supplemental Figures for "Revisiting a GWAS peak in *Arabidopsis thaliana* reveals possible confounding by genetic heterogeneity"

**Contents**

Fig. S1 GWAS peaks on *FLC* expression.

Fig. S2 Sequences of *NDX1* (Col-0 and 6180) with the domain information.

Fig. S3 Phenotypes of mutant lines.

Fig. S4 Extent of linkage disequilibrium.

Fig. S5 Conditional association of the SNP on *AOP2* with flowering time and *FLC* expression.

Fig. S6 Conditional association of the chr4 region with *FLC* expression.

Fig. S7 Heterogeneous haplotype structure of the chr4 region including *GA1* and *NDX1.*

Fig. S8 Linkage disequilibrium distribution of chromosome 4.

Fig. S9 GWAS and conditional association of the chr4 region with *FLC* expression and flowering time in global populations.

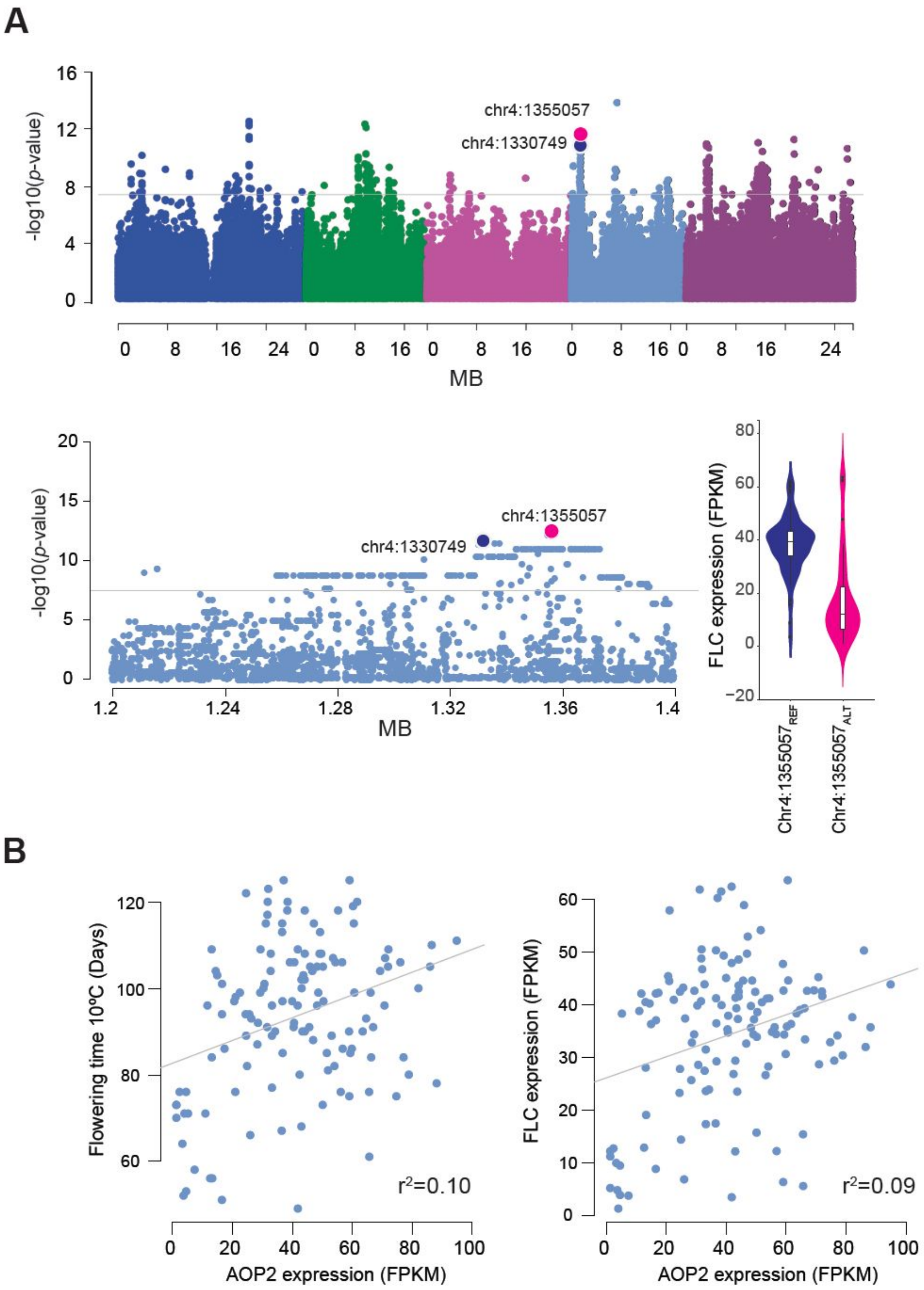


**Fig. S1 GWAS peaks on *FLC* expression.** (A) Genome-wide Manhattan plot for *FLC* expression at 10°C in 132 Swedish lines, with SNPs reported in Atwell et al. (2010) highlighted in blue (chr4: 1330749), and the strongest association in magenta (chr4: 1355057) with zoom-in on the peak. Violin plot showing the difference in *FLC* expression between lines carrying major and minor alleles at chr4:1355057. (B) Correlation of *AOP2* expression with flowering time (left) and *FLC* expression (right) in the same population ($n$=132). Grey lines are regression lines.

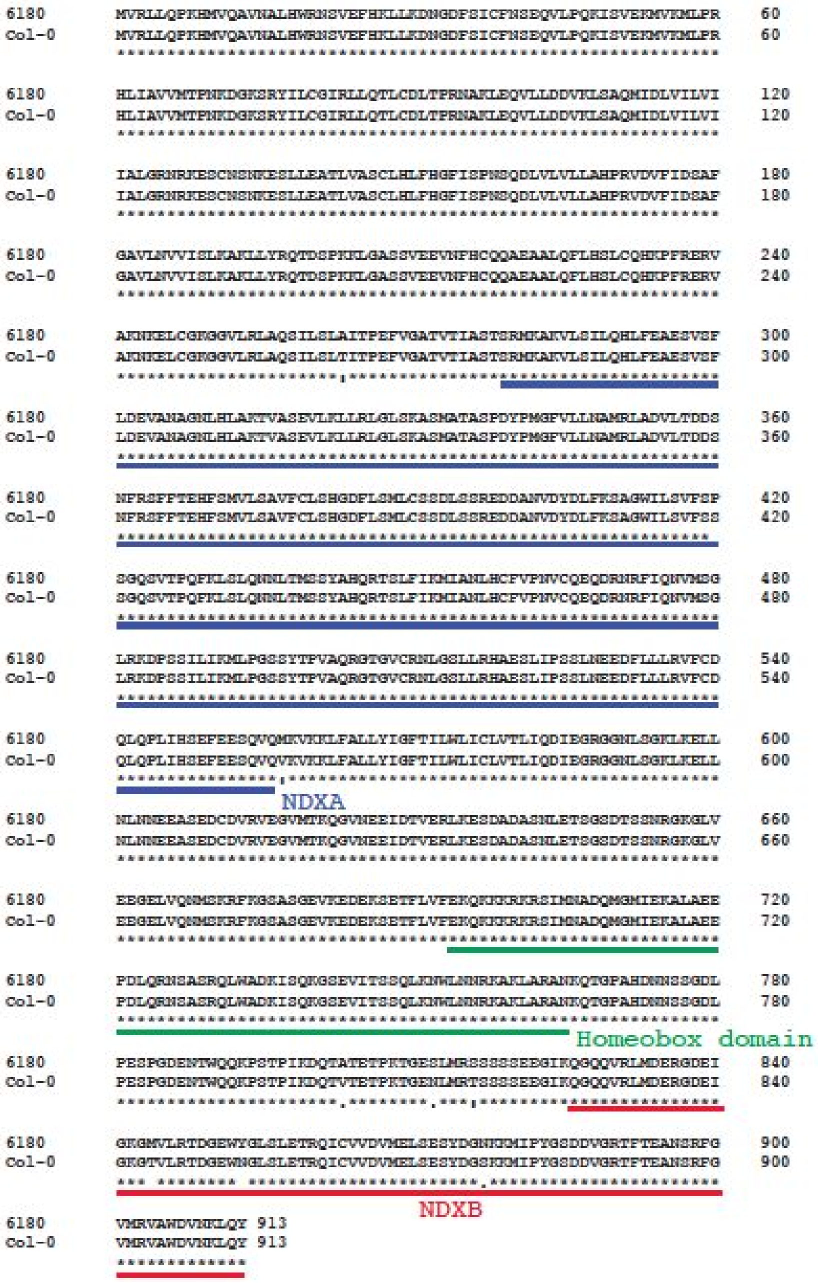


**Fig. S2 Sequences of *NDX1* (Col-0 and 6180) with the domain information.** Amino acid sequences of TÄL 07 (line 6180 carrying the minor allele of chr4: 1355057b) and Col-0 predicted by Sanger sequences. Underlined conserved domains NDX-A, Homeobox, and NDX-B were annotated according to Sun et al. (2013).

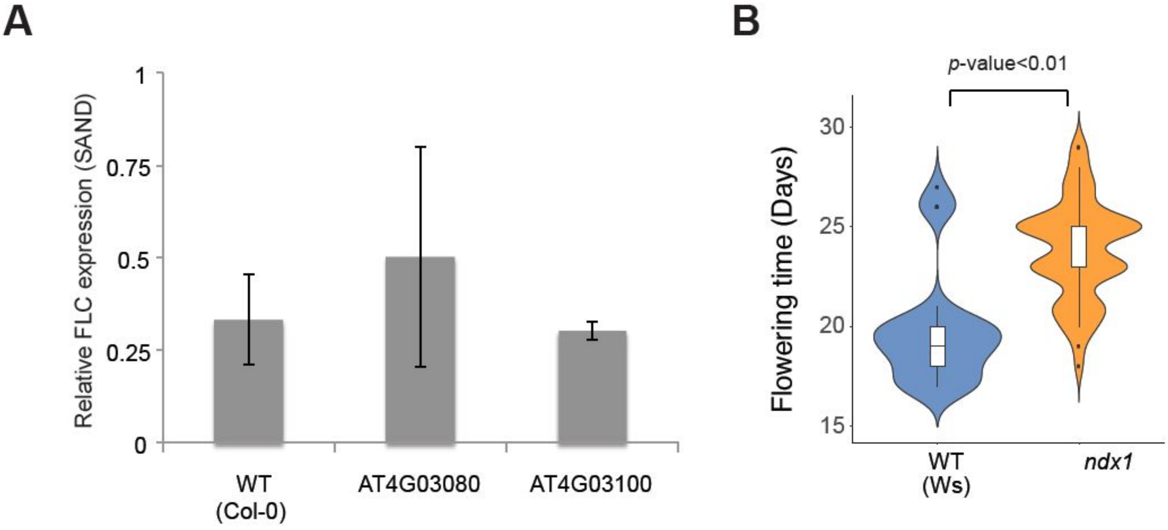


**Fig. S3 Phenotypes of mutant lines.** (A) *FLC* expression in loss-of function mutants near by *NDX1*. Error bars show standard deviation. There were no significant differences of *FLC* expression between WT (Col-0) and mutants (two-tailed Welch's *t*-test). (B) Violin plot showing flowering time of *ndx1.* Horizontal lines in the box plots show median. *P*-value was calculated by two-tailed Welch's t-test.

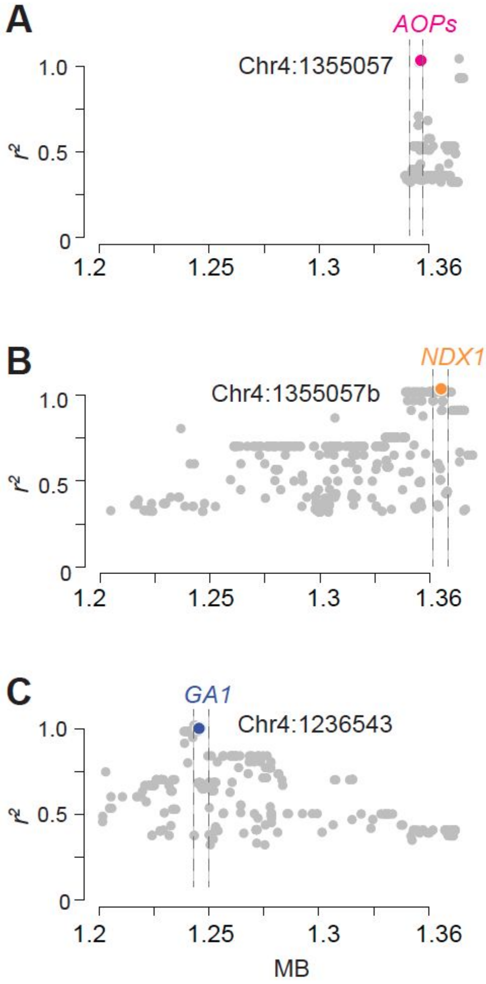


**Fig. S4 Extent of linkage disequilibrium.** Linkage disequilibrium ($r^2$) from the original *AOP2* SNP (A), the *NDX1* haplotype (B), and the *GA1* SNP (C). Vertical dash lines indicate the coding region of the candidate genes (+/- 3kb). $r^2$ for *NDX1* haplotype was calculated using chr4:1355057b genotypes (see Fig. 3). Only SNPs with $r^2$>0.25 were plotted.

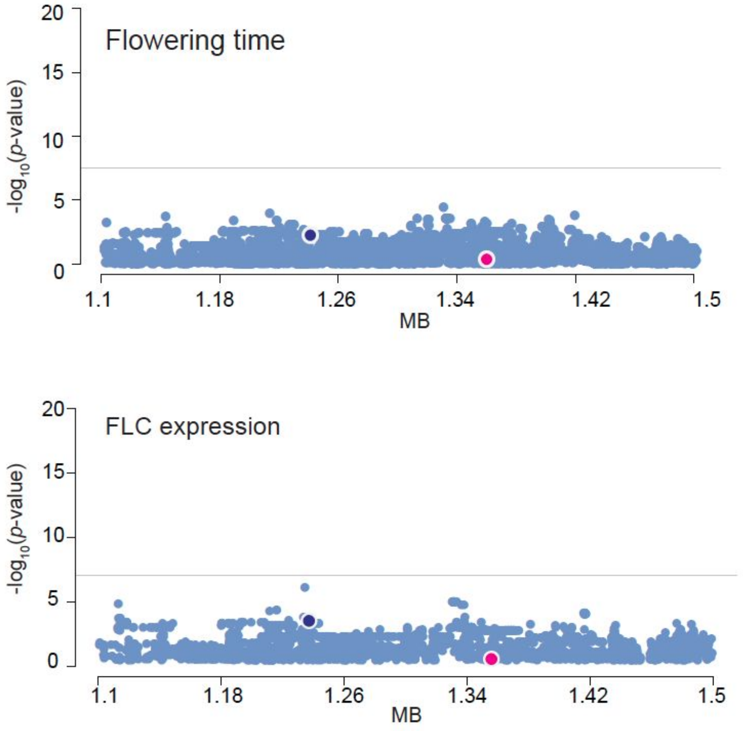


**Fig. S5 Conditional association of the SNP on *AOP2* with flowering time and *FLC* expression.** Zoom-in Manhattan plots of cofactor analysis of flowering time and *FLC* expression. For flowering time (top) and *FLC* expression (bottom), chr4:1355057 were used as the cofactor. Chr4:1355057 and chr4:1236543 were shown in magenta and blue, respectively. Horizontal lines show Bonferroni threshold (*p*-value=0.05).

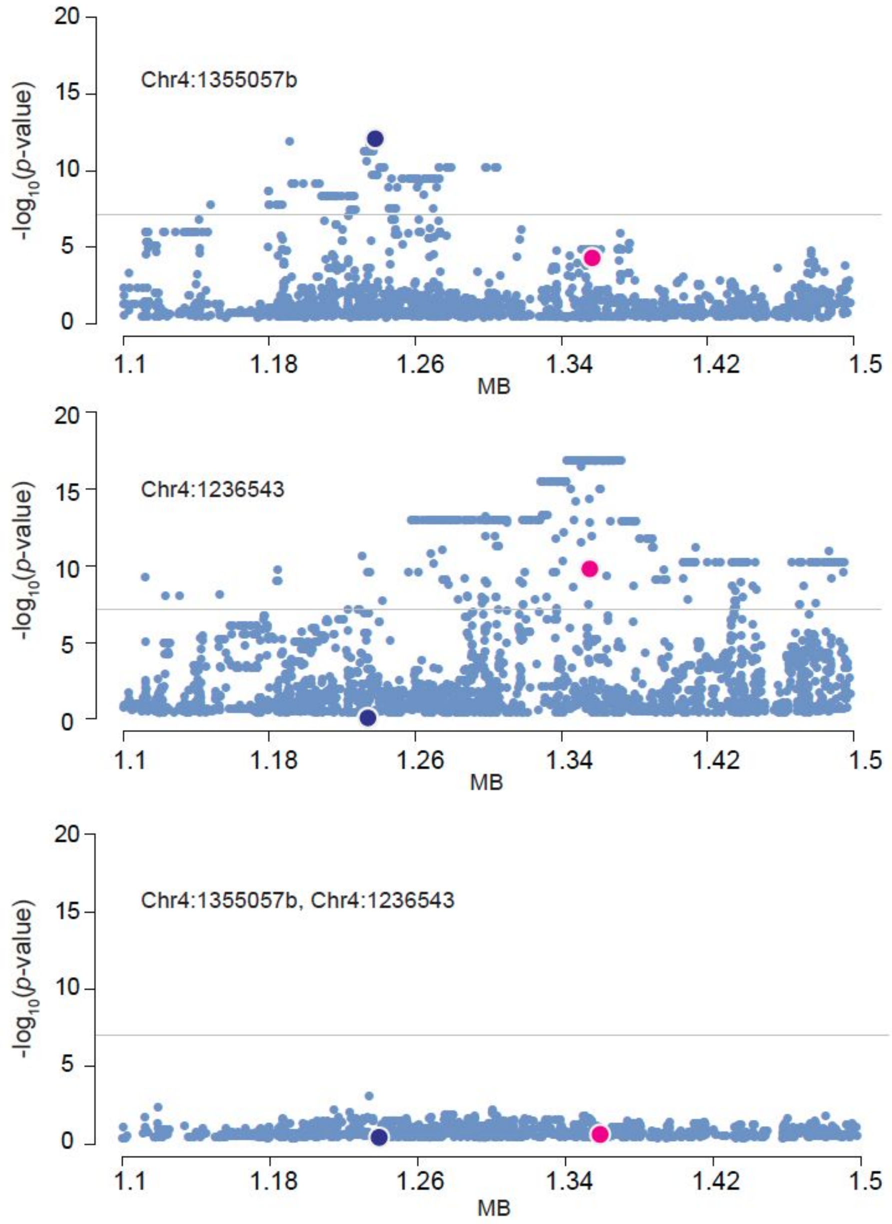


**Fig. S6 Conditional association of the chr4 region with *FLC* expression.** Zoom-in Manhattan plots of cofactor analysis of *FLC* expression. Chr4:1355057b (top), chr4:1236543 (second), and chr4:1355057b and 1236543 (bottom) were used for the cofactors, respectively. Chr4:1355057 and chr4:1236543 were shown in magenta and blue, respectively. Horizontal lines show Bonferroni threshold (*p*-value=0.05).

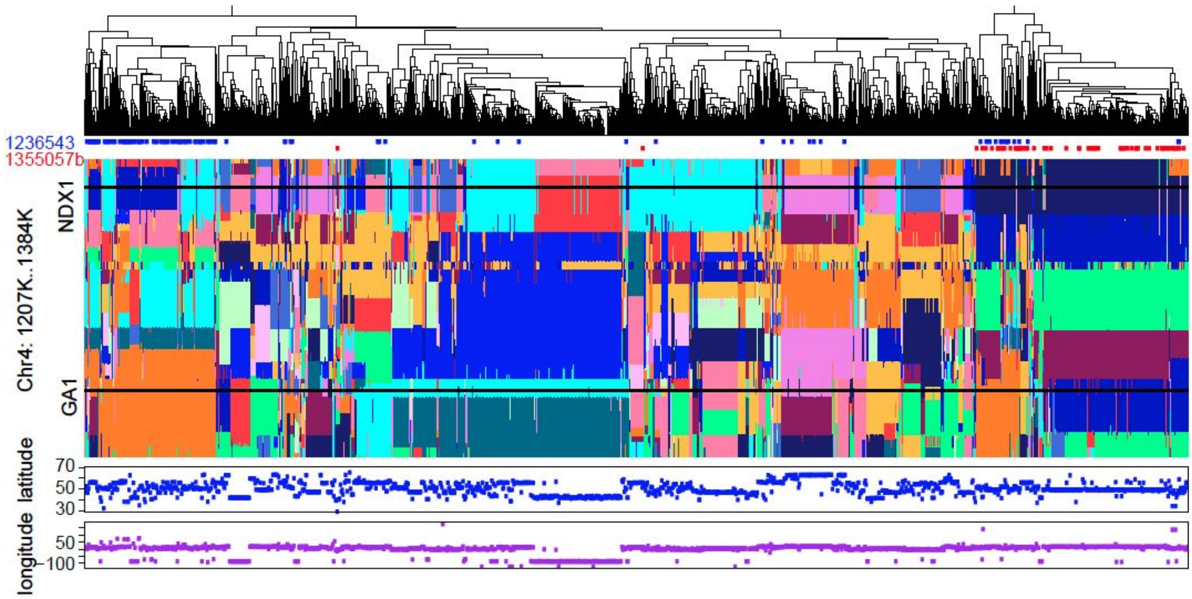


**Fig. S7 Heterogeneous haplotype structure of the chr4 region including *GA1* and *NDX1.*** Haplotype structure (chr4:1206543-1385057) in 1307 global *A. thaliana* lines (Horton *et al.*, 2012) was analyzed as described in Li et al. (2016). The top panel is a phylogenetic tree based on the SNPs in the analyzed region. Tips of the tree indicate minor alleles of chr4:1236543 and chr4:1355057b. The middle panel shows the genetic structure. Each line is represented in a column, and each SNP in the 180 Kbp window is represented in a row. At each SNP position, colors indicate the most likely haplotype membership for each line, as determined by fastPHASE analysis. Vertical black lines indicate locations of *GA1* (chr4:1236543) and *NDX1* (chr4:1366762 as a proxy of chr4:1355057b). The bottom panel shows the latitude and longitude of the original place that the lines were collected.

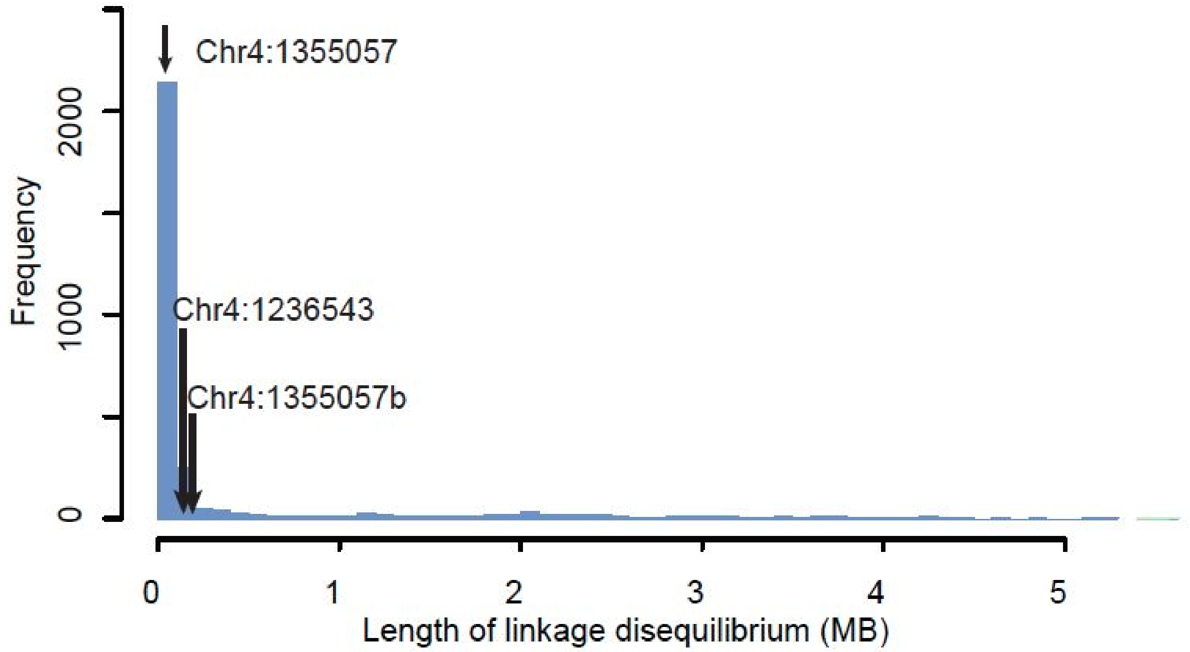


**Fig. S8 Linkage disequilibrium distribution of chromosome 4.** Length of linkage disequilibrium ($r^2$=0.6) from randomly chosen 3,000 SNPs on chromosome 4. Black arrows show the length from chr4:1355057, chr4:1355057b, and chr4:1236543.

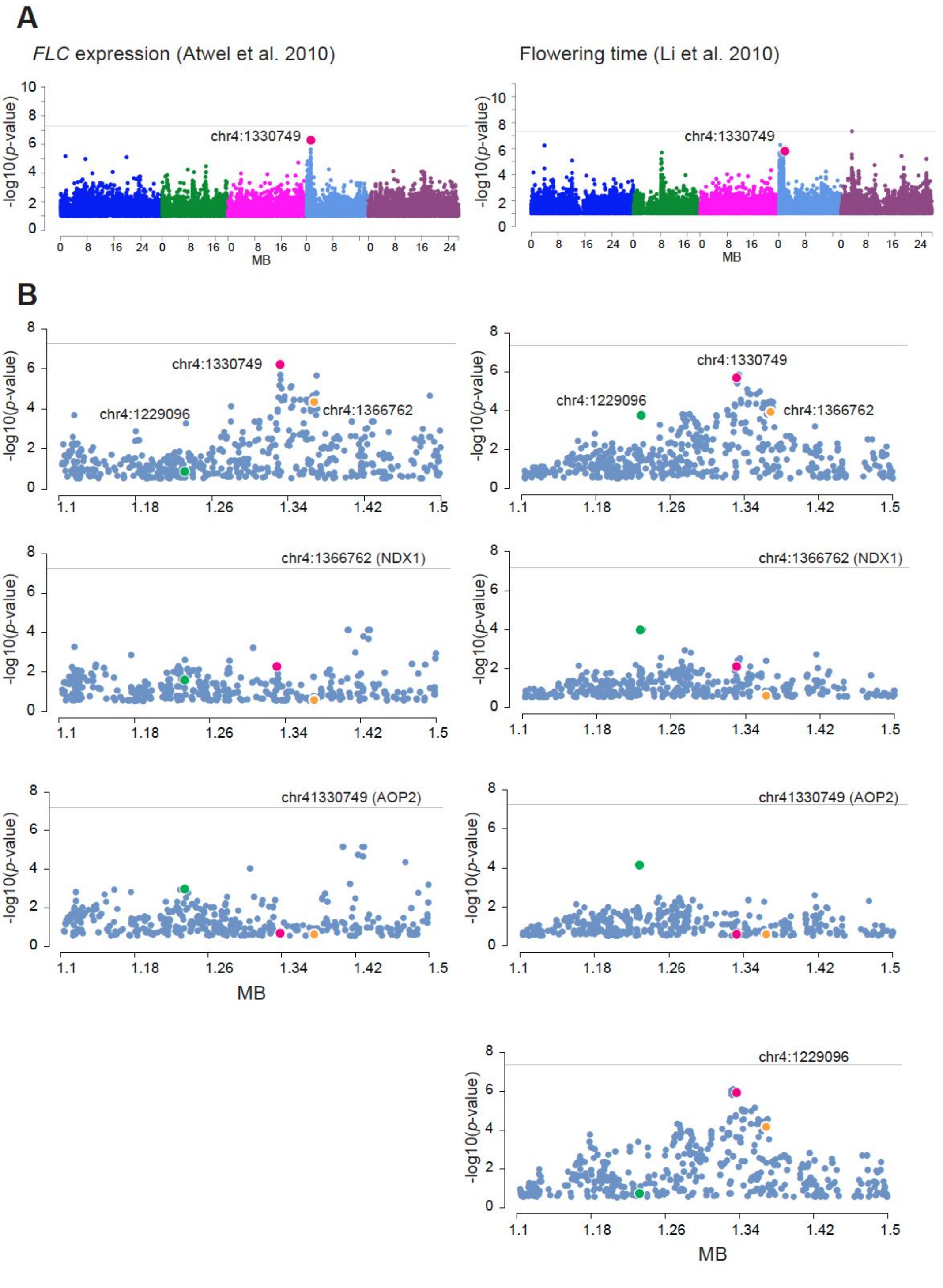


**Fig. S9 GWAS and conditional association of the chr4 region with *FLC* expression and flowering time in global populations.** (A) *FLC* expression (Atwel et al. 2010; *n*=167) and flowering time in Sweden spring condition in 2008 (Li et al. 2010; *n*=349) were reanalyzed by LMM using the original 250K SNP chip genotypes (Horton *et al.*, 2012). (B) Zoom-in Manhattan plots of cofactor analysis. Chr4:1366762 (top), chr4:1330749 (second), and chr4:1229096 (bottom) were used for the cofactors, respectively. Chr4:1366762, chr4:1330749, and chr4:1229096 were shown in orange, pink and green, respectively. Chr4:1366762 is used as a proxy of 1355057b ($r^2$ between 1355057b and 1366762 is 1 in the Swedish population). Chr4:1229096 is identified by GWAS for flowering time (Li et al. 2010) as the highest peak near by chr4:1236543. Horizontal lines show Bonferroni threshold (*p*-value=0.05).
